# Microbial-Derived Exerkines Prevent Skeletal Muscle Atrophy

**DOI:** 10.1101/2024.05.29.596432

**Authors:** Taylor R. Valentino, Benjamin I. Burke, Gyumin Kang, Jensen Goh, Cory M. Dungan, Ahmed Ismaeel, C. Brooks Mobley, Michael D. Flythe, Yuan Wen, John J. McCarthy

## Abstract

Regular exercise yields a multitude of systemic benefits, many of which may be mediated through the gut microbiome. Here, we report that cecal microbial transplants (CMTs) from exercise-trained vs. sedentary mice have modest benefits in reducing skeletal muscle atrophy using a mouse model of unilaterally hindlimb-immobilization. Direct administration of top microbial-derived exerkines from an exercise-trained gut microbiome preserved muscle function and prevented skeletal muscle atrophy.

## Main

The human gut microbiome is a collection of trillions of microorganisms inhabiting the gastrointestinal tract. These microorganisms act as a dynamic extension of the human host, serving many functions essential to our well-being^1^, and dysregulation of the microbiome can severely influence human health^2-6^. Recent evidence has tied the gut microbiome and microbial-derived metabolites to phenotypic adaptations in skeletal and cardiac muscle^7-11^. Notably, we reported that dysbiosis of the gut microbiome results in maladaptation in skeletal muscle in response to exercise-training^7^, demonstrating the importance of the gut microbiome in skeletal muscle adaptation. Given the relationship between the gut microbiome and exercise adaptation, efforts have been dedicated to determine if the benefits of exercise are able to be conferred via the gut microbiome^12^.

In the present study, we interrogated whether gut microbiome manipulations are able to ameliorate skeletal muscle atrophy during disuse. Cecal microbial transplants (CMTs) were utilized to transfer the gut microbiomes from sedentary or exercised donors into recipient mice (SED and EXR, respectively) for 5 weeks before recipient mice underwent unilateral hindlimb immobilization (HLI) via casting for 10 days to induce skeletal muscle atrophy (Fig. 1A). We focused our analysis on the soleus muscle because it undergoes the greatest magnitude atrophy of hind limb muscles with disuse. There was a significant decrease in the mean fiber cross-sectional area (CSA) in the soleus muscle of the casted leg compared to the control (i.e., free) leg in the SED and EXR groups (Fig. 1B), indicating atrophy in both groups. However, there was a significantly greater decrease in the soleus mean fiber CSA of the casted leg relative to the control leg in the SED group compared to the EXR group, a result that was not fiber-type specific (Fig. 1C). These data indicate that skeletal muscle atrophy induced via casting is ameliorated by the transplants of the gut microbiome from exercised donors.

**Figure 1:**
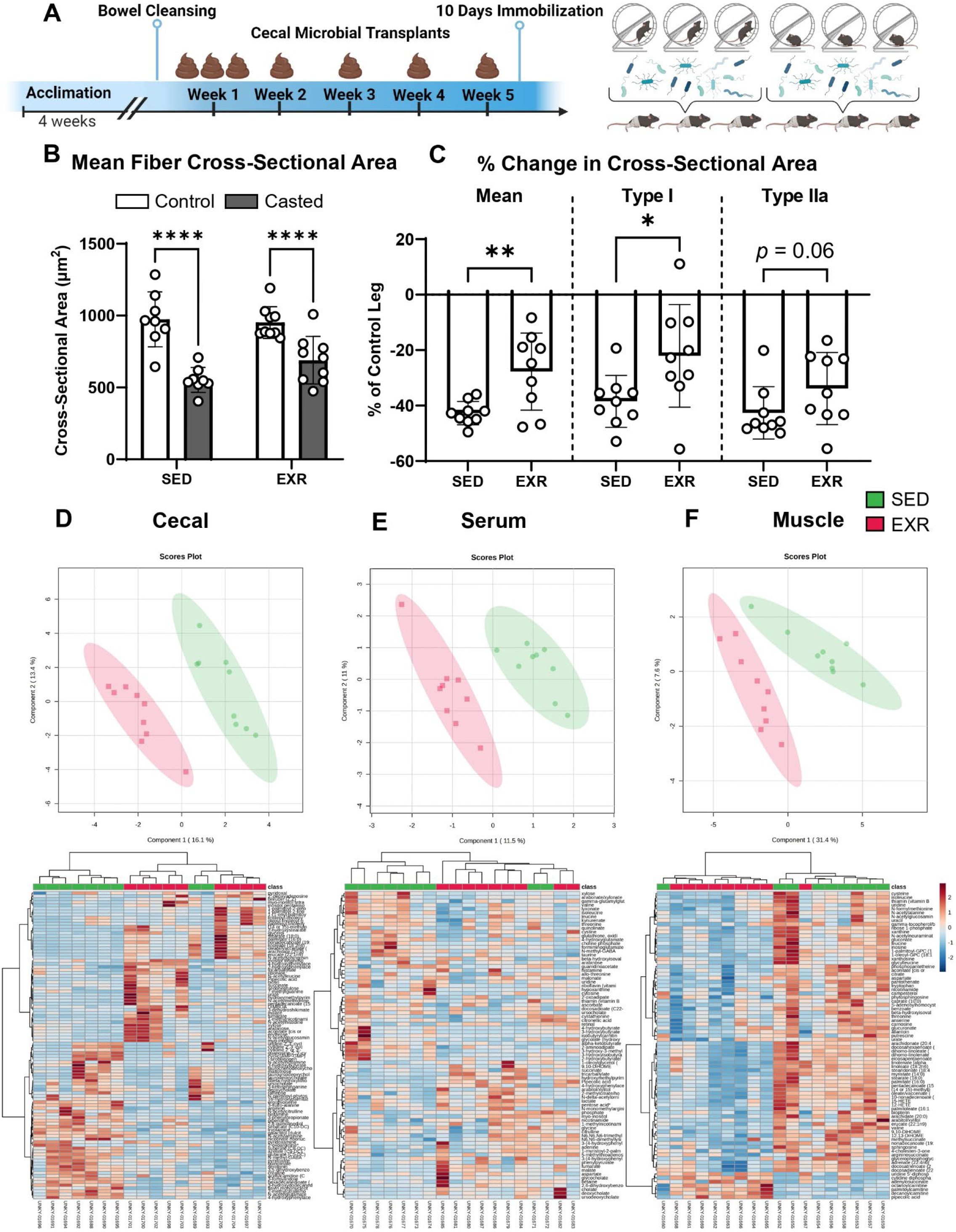
Microbial Transplants from Exercised Trained Donors Ameliorate Skeletal Muscle Atrophy. Female C57BL/6J mice (n = 9) received cecal microbial transplants from sedentary (SED) or exercised (EXR) donors for five weeks, followed by ten days of unilateral hindlimb immobilization via casting. Cecal content was pooled from three donors before transfer into recipient mice (A). Atrophy was assessed as the differences in mean cross-sectional area of the soleus between the control and casted leg in the SED and EXR groups (B). The magnitude of atrophy is expressed as the percent change in mean, type I, and type IIa cross-sectional area of the casted leg relative to the control leg in both groups (C). Metabolomics were performed on the cecal content (D), serum (E), and skeletal muscle (F); differences in the metabolomic profile of the mice receiving cecal microbial transplants from SED and EXR donors are expressed via Partial Least-Squares Discriminant Analysis (PLS-DA) plots and Hierarchical Cluster Analysis (HCA) heatmaps for each sample type. * = *p* < 0.05; ** = *p* < 0.01; **** = *p* < 0.0001.

In order to characterize the metabolomic response to CMTs, we performed untargeted metabolomics on the cecal content, serum, and gastrocnemius muscle from the SED and EXR groups. The metabolomic profiles in the cecum, serum, and skeletal muscle between the SED and EXR groups displayed clear differences as observed by Partial Least-Squares Discriminant Analysis (PLS-DA) plots and Hierarchical Cluster Analysis (HCA) heatmaps (Fig. 1D-F). These data demonstrate that transplantation of the gut microbiome from exercise-trained donors is capable of altering local and peripheral tissues within the recipients, despite the recipients remaining sedentary. The metabolomic outputs for each sample type (i.e., cecal content, serum, and skeletal muscle) were integrated and mean decrease accuracy (MDA) analyses were performed via random forest analysis in order to determine the predictive accuracy and metabolite importance among provided candidate features. Pipecolic acid, a lysine metabolite, and succinate, a Kreb’s cycle intermediate, were identified as the top two microbial-derived exerkines (MDE) predictive of the EXR group (Fig. 2A). These MDEs were administered via hydrogel separately (PIP and SUC, respectively) or together (PAS) for 10 days during which mice underwent HLI to induce skeletal muscle atrophy (Fig. 2B). Soleus mean fiber CSA decreased in the casted leg with respect to the control leg in the vehicle (VEH), SUC, and PIP groups; however, PAS maintained the CSA of the casted leg (Fig. 2C), an affect which was not fiber type specific (Fig. 2D-E). The total area of the muscle cross-section was also maintained with PAS treatment (Fig. 2F). Moreover, the maximal torque produced in the casted leg was maintained relative to the control leg in the PAS group, suggesting that the maintenance of muscle size resulted in a preservation of muscle function (Fig. 2G). Collectively, these data show that MDEs from an exercise-trained gut microbiome are able to prevent skeletal muscle atrophy and preserve muscle function upon direct administration to mice undergoing HLI.

**Figure 2:**
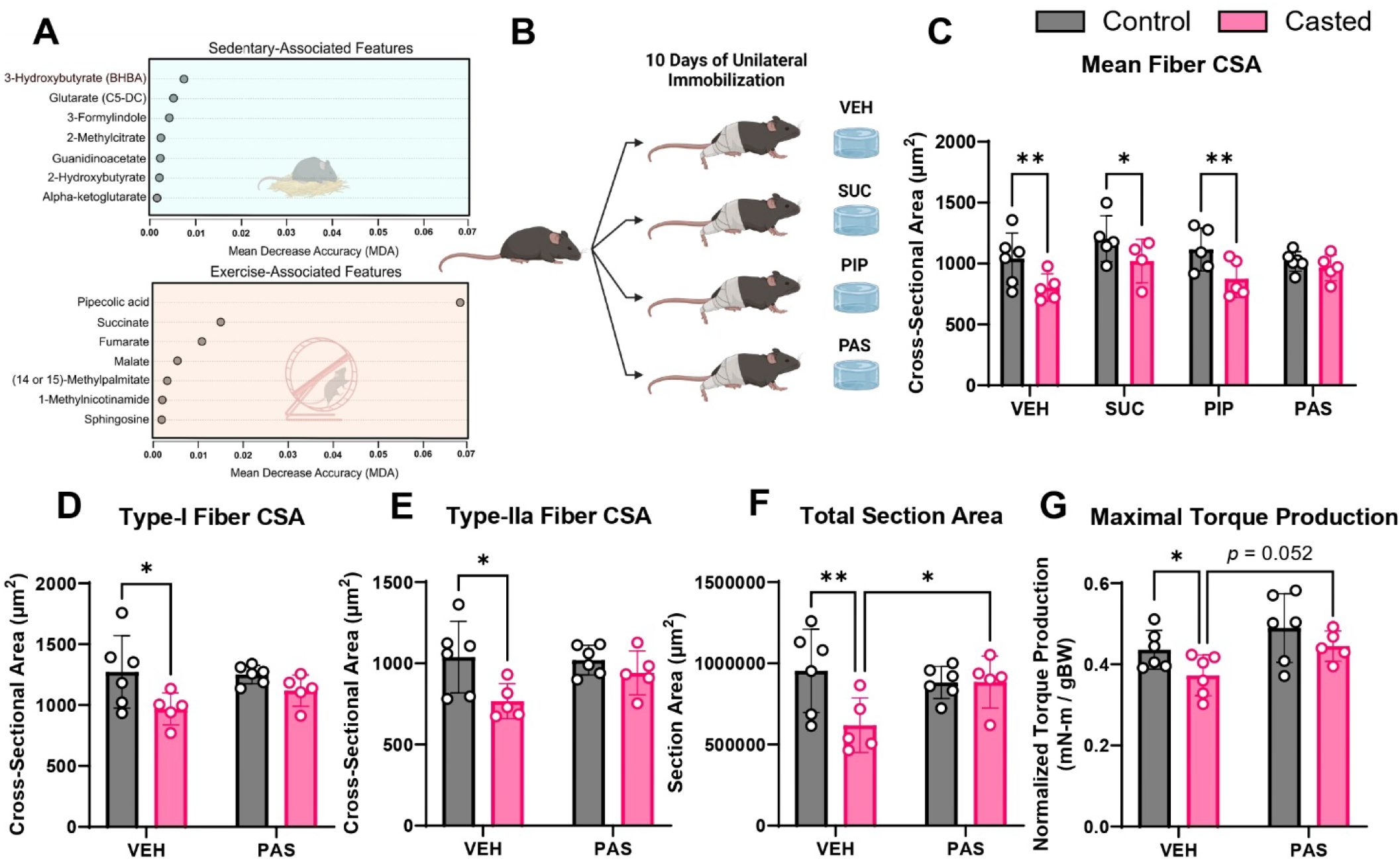
Microbial-Derived Exerkines Pipecolic Acid and Succinate Prevent Skeletal Muscle Atrophy and Preserve Muscle Function. Metabolomic analyses revealed pipecolic acid and succinate as the top microbial-derived exerkines derived from an exercise-trained gut microbiome (A). Female C57BL/6J mice (n = 5-6) were administered vehicle (VEH), succinate (SUC; 85.0 ± 7.7 mg/kg/d), pipecolic acid (PA; 170.0 ± 15.4 mg/kg/d), or pipecolic acid and succinate (PAS) through a hydration gel during ten days of unilateral hindlimb immobilization (B). Atrophy was assessed as the differences in mean (C), type I (D), and type IIa (E) cross-sectional area (CSA), as well as total section area (F), of the soleus between the control and casted leg. Muscle function was assessed as maximal torque production (G). * = *p* < 0.05; ** = *p* < 0.01.

In conclusion, the present study demonstrates that microbiome transplants from exercised-trained donors are able to ameliorate skeletal muscle disuse atrophy. Moreover, the administration of MDEs from an exercise-trained microbiome was able to reproduce this preservation of skeletal muscle mass along with the maintenance of muscle function. Previous groups have shown that specific microbes and/or microbial-derived products are able to affect skeletal muscle size and function^13-15^. However, this work leveraged exercise training to alter the composition and function of the gut microbiome^16,17^ in order to assess the therapeutic efficacy of an exercise-trained microbiome and its associated products. Other seminal papers have demonstrated the efficacy and therapeutic potential of an exercise-trained gut microbiome^12^.

The present study has built upon this work by demonstrating the ability of the gut microbiome to elicit therapeutic benefits on target tissues directly. These results demonstrate for the first time that an exercise-trained microbiome and associated metabolites elicit phenotypic effects on adult skeletal muscle corresponding with typical exercise adaptation (i.e., the maintenance of muscle mass during disuse as a result of exercise preconditioning^18^). Further, these findings serve as a proof of concept that some of the benefits of regular exercise are, in part, mediated by the gut microbiome. Given the widespread prescription of exercise as a therapeutic intervention for conditions such as cardiovascular disease, Alzheimer’s disease, aging, diabetes, etc., the findings of the present study provide compelling evidence to support leveraging an exercise-trained microbiome to treat various diseases positively impacted by exercise. Future efforts will be directed towards determining how MDE treatment is able to ameliorate disuse atrophy and establishing the efficacy of an exercise-trained gut microbiome to treat mouse models of Alzheimer’s Disease and aging.

## Methods

### Mice

12-week-old wild-type female C57BL/6J mice were purchased from The Jackson Laboratory and allowed to acclimate to the university vivarium for >4 weeks to allow the gut microbiome to stabilize^19^. For experiments involving the gut microbiome transfers, mice were randomly assigned to one of the following groups: sedentary donor, exercised donor, recipient from sedentary donor (SED), or recipient from exercised donor (EXR). During metabolite administration experiments, mice were randomly assigned to receive vehicle (VEH), succinate (SUC), pipecolic acid (PIP), or a combination of both succinate and pipecolic acid (PAS).

### Exercise Training

Donor mice were subjected to 8 weeks of progressive weighted wheel running (PoWeR) as described previously^20^. Briefly, mice were singly housed in running wheel cages with free access to the running wheel for one week of acclimation. After acclimation, weights consisting of 2g in week 1, 3g in week 2, 4g in week 3, 5g in weeks 4 and 5, and 6g in weeks 6-8 were added to one side of the wheel. Sedentary mice were singly housed under the same conditions with a locked running wheel.

### Cecal Microbial Transplant

The transplantation of cecal contents to a new host has recently emerged as a successful method of transplanting the gut microbiome in murine models^21,22^. Cecal contents were collected from exercised and sedentary donor mice and resuspended into sterile PBS at a concentration of 200mg/ml, which were stored -80°C. Prior to the first transfer, recipient mice were treated with polyethylene glycol (PEG) (SLBZ3934; Sigma-Aldrich, St. Louis, MO) to clear the existing microbiome^23^. After the mice received boluses of 200μL of 425g/L PEG via oral gavage every 20 minutes for 1 hour (4 total), they were placed into empty, sterile cages for 6 hours prior to the first transfer. Cecal contents collected from three exercise-trained or sedentary mice were pooled and transferred via oral gavage (200μL per transfer) to three anesthetized EXR and SED recipient mice, respectively. Recipient mice received transfers on three consecutive days, followed by a single transfer per week for the next 4 weeks, resulting in 7 total transfers over the course of 5 weeks. (Fig. 1A). Following the final transfer, mice were unilaterally casted to induce skeletal muscle atrophy for 10 days (see below).

### Unilateral Hindlimb Immobilization

Recently, a model of hindlimb immobilization utilizing casting was developed^24^ and subsequently modified by our lab using 3-D printing technology. This model allows the contralateral leg to remain mobile and serve as a control, thus limiting inter-animal variation in muscle size. Briefly, the leg is secured via Velcro into a 3-D printed mold in full extension at the knee and plantarflexion at the ankle, then covered with an external cast to prevent escape. Mice were casted for 10 days in all immobilization experiments.

### Immunohistochemistry

Following immobilization, hindlimb musculature (soleus and gastrocnemius) from both lower hindlimbs was carefully excised, weighed, and either covered in Tissue-Tek O.C.T. Compound (4583; Sakura Finetek, Torrance, CA) and snap-frozen in liquid nitrogen-cooled isopentane in preparation for immunohistochemistry (IHC) (soleus), or directly snap-frozen in liquid nitrogen for use in metabolomic analyses (gastrocnemius, see *Metabolomics* section below). Fiber-type staining procedures were carried out as previously described^25^. Briefly, soleus samples were cryosectioned at -24°C to obtain 7μm sections and dried for at least 1 hour before staining.

Unfixed sections were incubated at room temperature for 90 minutes with antibodies against myosin heavy chain types I (BA.D5), IIA (SC.71), and IIB (BF.F3) (1:100; Developmental Studies Hybridoma Bank, Iowa City, IA), as well as rabbit anti-laminin IgG (1:100; L9393; Sigma-Aldrich). The samples were then incubated with fluorescence-conjugated secondary antibodies against the various mouse immunoglobulin subtypes for 1 hour. Muscle sections were imaged using either a Zeiss upright fluorescent microscope (Zeiss AxioImager M1 Oberkochen, Germany) or an Olympus BX61VS Upright Fluorescent Microscope (Evident Scientific, Bethlehem, PA) at 20x magnification. MyoVision, an unbiased automated image analysis program developed by our lab, was used to determine myofiber CSA and myonuclear abundance in a blinded manner^26^.

### Metabolomics

Cecal contents, serum, and gastrocnemius samples were collected from SED and EXR groups at sacrifice. For the cecal contents, the cecum was removed and on a sterile dish the contents were carefully removed and collected. For serum, blood was collected at the point of sacrifice, allowed to clot and centrifuged at 150 x g for 15 minutes. The supernatant was removed and centrifuged again at 300 x g for 15 minutes. The supernatant was collected and stored at -80°C for further analysis. Cecal contents and gastrocnemius muscles were collected, immediately snap frozen in liquid nitrogen and stored at -80°C. Samples were sent to Metabolon, Inc. (Morrisville, NC) for untargeted metabolite profiling.

### Metabolomic Data Analysis

In the discriminative analytical approach for SED and EXR using Partial Least-Squares Discriminant Analyses (PLS-DA), the explanatory power (R^2^) and predictive ability (Q^2^) of each model were assessed. For a perfect model, both R^2^ and Q^2^ values are defined as 1.0 (100%). Moreover, Hierarchical Cluster Analyses (HCA) were performed using the top 80 metabolites in each sample type (i.e., cecal content, serum, and skeletal muscle) in order to observe the metabolic discriminative patterns between SED and EXR. In order to identify important biomarkers distinguishing SED and EXR, metabolomic profiles from the three sample types were integrated and MDA analyses, a widely used omics analysis method^27-29^, were conducted using random forest analysis to determine predictive accuracy and the importance of metabolic features among the given candidates. All data processing and analyses were performed using the online software MetaboAnalyst 6.0^30^.

### Microbial-Derived Exerkine Administration

For experiments involving the direct administration of microbial-derived metabolites, herein known as microbial-derived exerkines (MDEs) mice received either vehicle, succinate (S7501; Sigma-Aldrich), pipecolic acid (P2519; Sigma-Aldrich), or both succinate and pipecolic acid via MediGel Sucralose hydrogels (74-02-5022; ClearH_2_O, Westbrook, ME). Mice received free access to the hydrogels for 10 days during hindlimb immobilization as described above. During this period, mice consumed an average of 85.0 ± 7.7 mg/kg/d and 170.0 ± 15.4 mg/kg/d of succinate and pipecolic acid, respectively. Following metabolite administration and immobilization, mice were sacrificed and the hindlimb muscles were prepared for IHC as described above.

### Statistics

Two-way ANOVAs were used to compare differences in muscle mass, mean CSA, fiber-type specific CSA, total section area, and maximal force production between control and casted legs within and between all groups. All statistical analyses were performed using GraphPad Prism version 10.0.3 for Windows (GraphPad Software, La Jolla, CA).

## Data Availability

All omic data sets generated during the course of this study will be made publicly available through Gene Expression Omnibus upon publication.

## Acknowledgements

This work was supported by the National Institutes of Health (R21AG071888 to J.J.M.).

